# Spindle assembly checkpoint-dependent mitotic delay is required for cell division in absence of centrosomes

**DOI:** 10.1101/2022.11.08.515699

**Authors:** KC Farrell, Jennifer T. Wang, Tim Stearns

## Abstract

The spindle assembly checkpoint (SAC) temporally regulates mitosis by preventing progression from metaphase to anaphase until all chromosomes are correctly attached to the mitotic spindle. Centrosomes refine the spatial organization of the mitotic spindle at the spindle poles. However, centrosome loss leads to elongated mitosis, suggesting that centrosomes also inform the temporal organization of mitosis in mammalian cells. Here we find that the mitotic delay in acentrosomal cells is enforced by the SAC in a MPS1-dependent manner, and that a SAC-dependent mitotic delay is required for bipolar cell division to occur in acentrosomal cells. Although acentrosomal cells become polyploid, polyploidy is not sufficient to cause dependency on a SAC-mediated delay to complete cell division. Rather, the division failure in absence of MPS1 activity results from mitotic exit occurring before acentrosomal spindles can become bipolar. Furthermore, prevention of centrosome separation suffices to make cell division reliant on a SAC-dependent mitotic delay. Thus, centrosomes and their definition of two spindle poles early in mitosis provide a “timely two-ness” that allows cell division to occur in absence of a SAC-dependent mitotic delay.

## Introduction

In eukaryotic cells, depolymerization of microtubules and the resultant mitotic arrest before anaphase led to the discovery of the ‘mitotic block’,^1,2^ now called the spindle assembly checkpoint (SAC). SAC activity prevents progression from metaphase to anaphase by inhibiting the anaphase-promoting complex/cyclosome (APC/C).^3–6^ The SAC, comprised of components largely enriched near kinetochores,^7–10^ becomes activated when defects in chromosome attachment are present.^7,8,11,12^ Thus, perturbance of spindle microtubules or certain microtubule motors leads to SAC activation.^7–9,13,14^ At the other end of each side of the spindle apparatus, centrosomes are attached to and focus the spindle poles in a dynein-dependent manner.^15,16^ Centrosomes are usually comprised of two centrioles and pericentriolar material (PCM) that contains many microtubule nucleation factors that stabilize microtubule minus-ends, and they are major points of microtubule minus-ends in the mitotic spindles of mammalian cells.^15,17–20^ However, in absence of centrosomes, cultured mammalian cells can still divide bipolarly.^21–23^ While centrosomes, when present, are major sites of microtubule nucleation, microtubules can be nucleated in mammalian cells by acentrosomal microtubule organizing centers (MTOCs)^24,25^ as well as the chromatin-mediated microtubule nucleation pathway.^26–28^ Nevertheless, absence of centrosomes alters bipolar spindle formation, such that spindles are monopolar early in mitosis and then resolve to become bipolar.^24,25,29^ The construction of such acentrosomal spindles occurs during a mitosis that is, on average, elongated.^30,31^ In this work, we tested whether SAC activity is necessary for this elongation, since the SAC is a well-conserved mechanism to delay mitotic progression. Furthermore, we asked whether the elongation is necessary for acentrosomal division to proceed and, if so, what centrosomes provide to negate the need for this elongation.

## Results and Discussion

### MPS1 activity is required for mitotic elongation and cell division in acentrosomal cells

We removed centrosomes to interrogate their contributions to the temporal progression of mitosis. Acentriolar, hTERT-immortalized RPE-1 cells were generated though 10d treatment with the PLK4 inhibitor centrinone (125 nM)^30^ or through deletion of *SASS6* which encodes a structural protein (SASS6) that templates centriole duplication.^32^ Acentriolar RPE-1 cells were generated in *TP53^-/-^* background, since RPE-1 cells with intact p53 pathways undergo G1 arrest in absence of centrioles.^30,31^ Acentriolar wild-type U2OS cells, which are already p16-deficient,^33^ were generated through 10d treatment with centrinone (125 nM). RPE-1 *TP53^-/-^* and U2OS cells treated with centrione showed significant increases in acentrosomal cells as compared to DMSO controls, as did *TP53^-/-^; SASS6^-/-^* cells in comparison to *TP53^-/-^*cells as measured by number of γ-tubulin and centrin-3 co-positive puncta, which respectively mark PCM and centrioles (Figure 1a-a’). RPE-1 *TP53^-/-^* and U2OS cells treated with centrinone as wells as *TP53^-/-^; SASS6^-/-^* cells will be referred to as “acentrosomal cells” for brevity. As has been previously reported,^30,31^ mitosis was elongated in acentrosomal cells compared to their appropriate controls (Figure 1b). However, the mitoses were not uniformly elongated during each stage. Rather, the time from nuclear envelope breakdown (NEBD) through anaphase onset was elongated in acentrosomal cells, while the duration of anaphase through cytokinesis was not significantly different compared to their appropriate controls (Figure 1b-b’).

**Figure 1.**
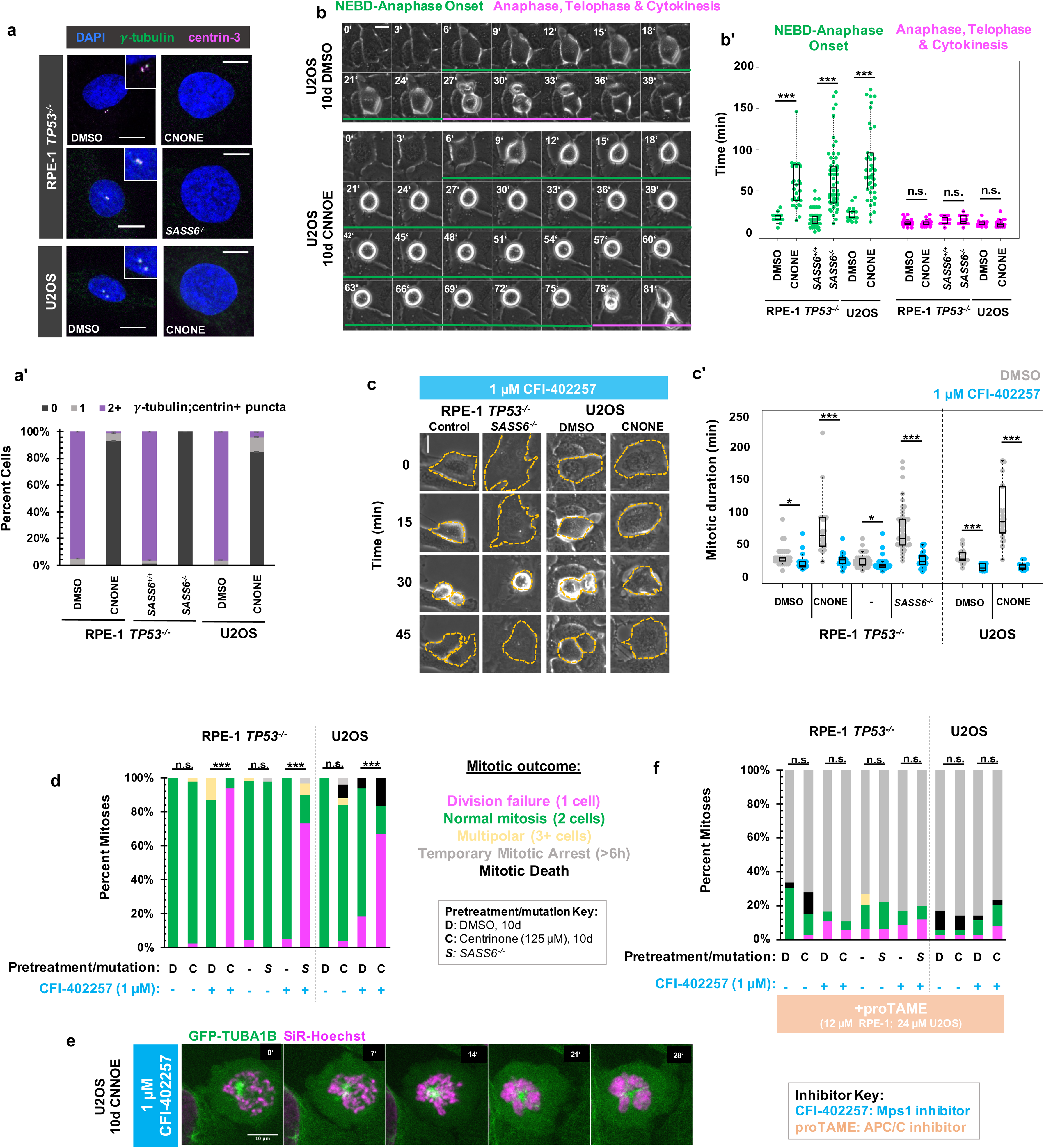
MPS1 activity is required for mitotic elongation and cell division in acentrosomal cells. **(a)** Immunofluorescence staining of control and acentriolar RPE-1 *TP53^-/-^* or U2OS cells. Acentriolar cells were created by treatment with 125 nM centrinone (CNONE) or by deletion of *SASS6*. DAPI is shown in blue, γ-tubulin in green, and centrin-3 in magenta. Insets are shows for cells in which centrosomes were present. **(a’)** Quantification of a. Graphed are means and S.E.M. Significance was determined through a Fisher’s exact test. *n*=100 cells per condition**. (b)** Live phase contrast imaging showing example mitosis in U2OS cells pre-treated with DMSO or centrinone (CNONE) for 10 days. Green bars below cells indicate the duration of nuclear envelope breakdown (NEBD) through the onset of anaphase, while magenta bars below cells indicate the duration of the completion of anaphase, telophase, and cytokinesis. **(b’)** Mitotic durations from NEBD through metaphase (green) and anaphase, telophase, and cytokinesis (magenta) as in (b). Time is in minutes. Points represent individual cells; boxplots represent mean and interquartile range. Significance was determined through Welch’s *t*-test. *n*>20 cells per condition. **(c)** Live phase contrast imaging showing example mitosis in U2OS cells pre-treated with DMSO or centrinone (CNONE) for 10 days before being imaging in 1 μM CFI-402257. **(c’)** Quantification of mitotic duration in cells of indicated genotype or treatment in DMSO (grey) or CFI-402257 (blue). Time is in minutes. Points represent individual cells; boxplots represent mean and interquartile range. Significance was determined through Welch’s *t*-test. *n*>25 cells per condition. **(d)** Quantification of daughter cell fate of cells of the given pre-treatment imaged in DMSO or CFI-402257 (1 μM). Shown are the percentages for each fate of the total mitotic observations. Significance was determined through a Fisher’s exact test. *n*>40 cells per condition. **(e)** Confocal timelapse imaging of U2OS cells pretreated for 10 d with 125 nm centrinone and imaged in 1 μM CFI-402257. Shown are endogenously tagged α-tubulin (*GFP-TUBA1B*) and DNA (Sir-Hoechst). **(f)** Quantification of daughter cells fates of cells of the given pre-treatment imaged in DMSO or CFI-402257 (1 μM) with concurrent treatment with proTAME (12 μM for RPE1 cells, 24 uM for U2OS cells). Shown are the percentages for each fate of the total mitotic observations. Significance was determined through a Fisher’s exact test. *n*>40 cells per condition. In all cases, not significant (n.s.) denotes *p*>0.05, * denotes p<0.05, **denotes p<0.01, and *** denotes p<0.001. All scale bars: 10 μm. **Figure1_Source_Data_1.zip: Source data for Figure 1**

The SAC is responsive to spindle defects that occur from NEBD through anaphase onset,^34^ which was the elongated mitotic time period in acentrosomal cells. Therefore, we tested whether mitotic elongation in acentrosomal cells depended on SAC activity. We inhibited MPS1 (<u>m</u>ulti<u>p</u>olar <u>s</u>pindle 1) kinase (also called Threonine Tyrosine Kinase, TTK), a necessary SAC component^35,36^ that begins high activity shortly before NEBD,^37^ via treatment with CFI-402257.^38,39^ Both control and acentriolar cells treated with CFI-402257 showed reduced mitotic durations (Figure 1c-c’) as measured by the time from visible NEBD to visible nuclear envelope reformation. Thus, MPS1 activity is necessary for the extended mitosis normally observed in acentrosomal cells.

While the majority of control cells treated with CFI-402257 underwent bipolar mitoses, the majority of acentriolar cells treated with CFI-402257 rounded up and spread back out, producing a single mononucleate daughter cell (Figure 1c,d). Acentriolar cells treated with a different MPS1 inhibitor, MPS1-IN-1^40^, also underwent mitotic events that produced only one daughter cell each (Figure 1-supplement 1). Examination of nuclear morphology revealed that the resultant cells were not binucleate, suggesting that the division failure was not a cytokinetic but a karyokinetic defect; however, the nuclei of acentrosomal cells after division in CFI-402257 were highly abnormal in morphology (Figure 1-Supplement 2).

To further examine mitosis in acentrosomal cells without MPS1 activity, we generated U2OS cells expressing N-terminally GFP-tagged α-tubulin (*GFP-TUBA1B*) from one allele of the *TUBA1B* locus, made using CRISPR/Cas9-mediated homologous recombination^41^ (Figure 1-supplement 3). Cells were treated with centrinone for 10 d prior to imaging and, for imaging, were incubated with a low concentration of SiR-Hoechst (50 nM) to stain DNA. In the mitotic cell shown (Figure 1e), a single α-tubulin punctum was present alongside condensed chromosomes. While the chromosomes eventually decondensed, a misshapen nucleus formed without cell division occurring.

The phenotypes observed, namely elongated mitosis and sensitivity of cell division to MPS1 inhibition, were reversible after 10 d of centrinone washout and return to normal centriole number (Figure 1-supplement 4). The cell division failure phenotype observed in acentriolar cells treated with CFI-402257 could be prevented via treatment with proTAME, ^42^ an inhibitor of the APC/C (Figure 1f, Figure S1b). ProTAME treatment resulted in a mitotic arrest of 6 h or more, regardless of centrosome status, indicating that the rounding up/sitting down behavior of acentrosomal cells treated with MPS1 inhibitor truly represented mitotic entry and exit in the absence of cell division.

### Delayed anaphase onset, rather than specific MPS1 activity, is necessary for successful acentrosomal cell division

We considered whether specifically the kinase activity of MPS1 was required for division in acentrosomal cells or rather its general role in promoting mitotic delay. Aurora B kinase becomes enriched at kinetochores and chromosomes during mitosis and regulates a variety of SAC and error-correction components.^10,43–45^ We therefore used inhibition of Aurora B to perturb the SAC function independent of MPS1 inhibition. To perturb Aurora B function, we incubated cells with 50 nM of the Aurora B inhibitor AZD1152, also known as Barasertib,^46^ and imaged cells live via phase microscopy (Figure 2a). Unlike with inhibition of MPS1, both control and acentrosomal cells underwent elongated cell divisions when treated with AZD1152 (Figure 2a’). Furthermore, in contrast to treatment with MPS1 inhibitors, both control and acentrosomal cells divided into two daughter cells the majority of the time (Figure 2a’’). Co-treatment with MPS1 and Aurora B inhibitors resulted in shortened mitosis in the acentrosomal cells and resulted in a similar division failure rate as with MPS1 inhibition alone (Figure 2a-a’’).

**Figure 2.**
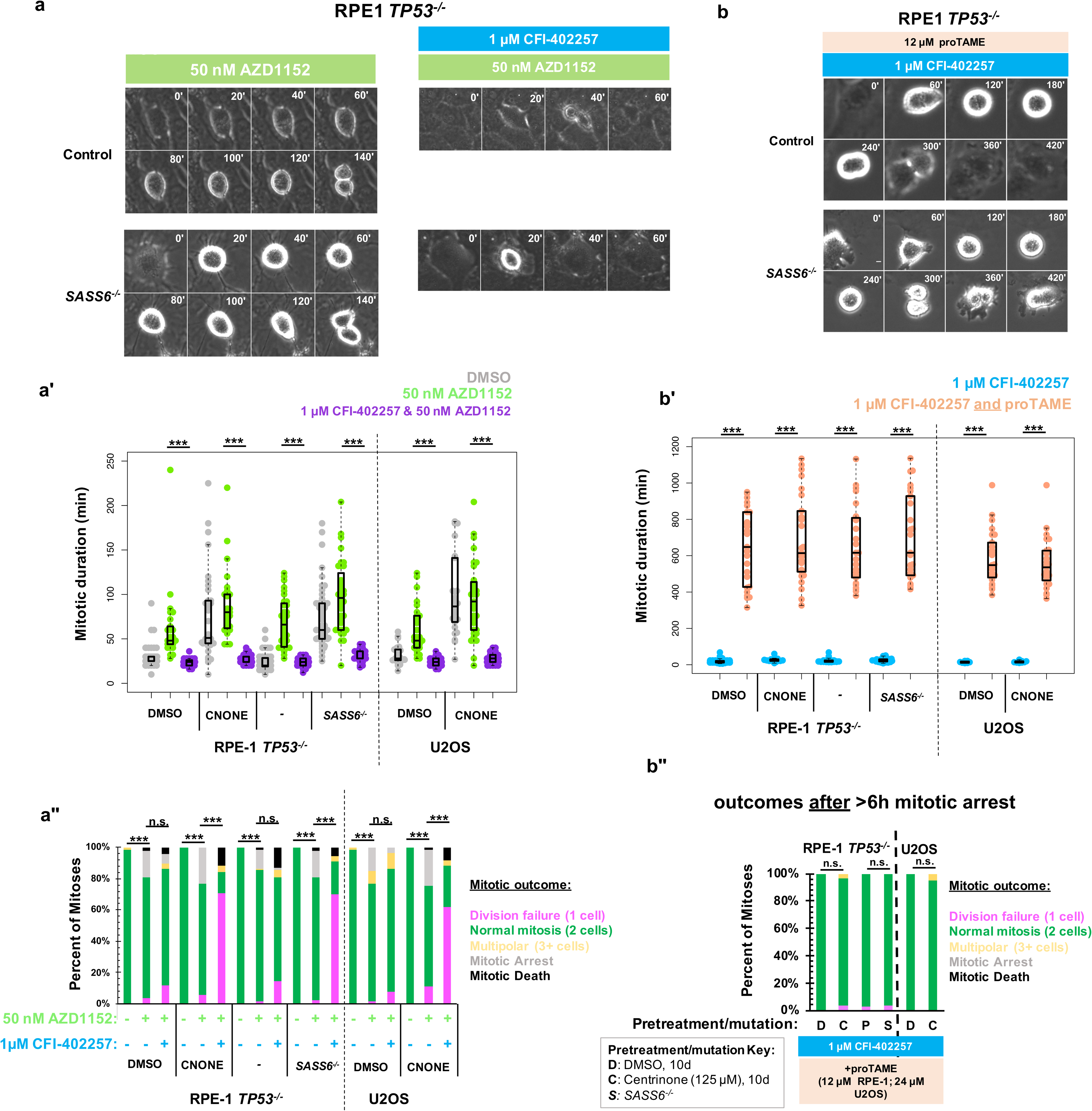
Delayed anaphase onset, rather than specific MPS1 activity, is necessary for successful acentrosomal cell division. **(a)** Example mitoses in RPE1 TP53^-/-^ or RPE1 *TP53^-/-^; SASS6^-/-^*cells treated with 50nM AZD1152 with or without 1μM CFI-402257. **(a’)** Quantification of mitotic duration in cells of indicated genotype or treatment in DMSO (grey), 50nM AZD1152 (green), or 50nM AZD1152 and 1μM CFI-402257 (purple). Points represent individual cells; boxplots represent mean and interquartile range. Significance was determined through Welch’s *t*-test. *n*>47 cells per condition. **(a’’)** Quantification of daughter number of cells of the given pre-treatment imaged in DMSO, 50nM AZD1152 or 50nM AZD1152 and 1μM CFI-402257. Shown are the percentages for each fate of the total mitotic observations. *n*>20 cells per condition. Significance was determined through a Fisher’s exact test. **(b)** Example mitoses in RPE1 TP53^-/-^ or RPE1 *TP53^-/-^; SASS6^-/-^* cells treated with 1μM CFI-402257 and proTAME (12 μM for RPE1 cells, 24 μM for U2OS cells). **(b’)** Quantification of mitotic duration in cells of indicated genotype or treatment in 1μM CFI-402257 (blue) or 1 μM CFI-402257 with 12 μM (RPE1 cells) or 24 μM (U2OS cells) proTAME. Points represent individual cells; boxplots represent mean and interquartile range. Significance was determined through Welch’s *t*-test. *n*>25 cells per condition. **(b’’)** Quantification of daughter number of cells of the given pre-treatment imaged in 1μM CFI-402257 (blue) or 1 μM CFI-402257 with 12 μM (RPE1 cells) or 24 μM (U2OS cells) proTAME after a mitotic arrest of greater than six hours. Shown are the percentages for each fate of the total mitotic observations. *n*>20 cells per condition. Significance was determined through a Fisher’s exact test. In all cases not significant (n.s.) denotes *p*>0.05 and *** denotes p<0.001. All scale bars: 10 μm. **Figure2_Source_Data_1.zip: Source data for Figure 2a, a’’,b, b’’**

Based on these results, we hypothesized that the mitotic delay caused by MPS1 activity was necessary for successful cell division in acentrosomal cells. We tested this by asking whether the phenotype of division failure in acentrosomal cells under MPS1 inhibition could be rescued by elongating mitosis independent of checkpoint activation. APC/C inhibition did result in increased mitotic arrest (>6 h) in cells regardless of MPS1 inhibition (Figure 1e). However, many of the acentrosomal cells treated with both proTAME and CFI-402257 eventually went on to divide, after the prolonged mitotic arrest. Treatment with both proTAME and CFI-402257 resulted in majority bipolar divisions in acentrosomal cells (Figure 2b-b”), in contrast to treatment with CFI-402257 alone (Figure 1d). This suggests that the division failure observed in acentrosomal cells treated with MPS1 inhibitor was due to mitotic shortening caused by preemptive anaphase onset and not other activities of MPS1.

### Polyploidy is not sufficient to confer MPS1-inhibition-sensitive division failure

We next asked what defect of an acentrosomal mitosis that requires a delay in anaphase onset to complete cell division. Acentrosomal cell divisions are more prone to chromosome missegregation and fragmentation,^31,47,48^ and populations of acentrosomal cells cultured for multiple generations are more likely to be aneuploid.^47^ We confirmed that our acentrosomal cells were polyploid by counting chromosome pairs in metaphase spreads (Figure 3a). A simple hypothesis would be that cell division in acentrosomal cells is sensitive to MPS1 inhibition due to a requirement for the SAC in cells with increased ploidy. In fact, previous results show that SAC activity was required for the mitotic elongation in tetraploid acentrosomal mouse embryos^49^ and that aneuploid cultured cells are more sensitive to SAC inhibition.^50^ We therefore tested the contribution of polyploidy to division failure during MPS1 inhibition. If polyploidy is sufficient for this phentotype in RPE1 and U2OS cells, then polyploid cells with centrosomes should also require a SAC-based delay to divide. We treated RPE1 *TP53^-/-^* cells with 50 μM blebbistatin, a myosin inhibitor,^51^ for 12 h to induce cytokinesis failure, which resulted in a mixed population of binucleate cells resulting from division failure and mononucleate cells that had not entered mitosis during the incubation and served as an internal control for normal ploidy (Figure 3b). We then removed the blebbistatin, treated with the MPS1 inhibitor CFI-402257, and assessed division outcomes. Under treatment with CFI-402257, mononucleate RPE1 *TP53^-/-^* cells divided into two daughters the majority of the time (Figure 3c-c’). Binucleate RPE1 *TP53^-/-^*cells divided the majority of the time, into either two or multiple (3+) daughters (Figure 3c-c’). Thus, polyploidy alone, as generated by cytokinesis failure, was not sufficient to cause division failure when MPS1 is inhibited.

**Figure 3.**
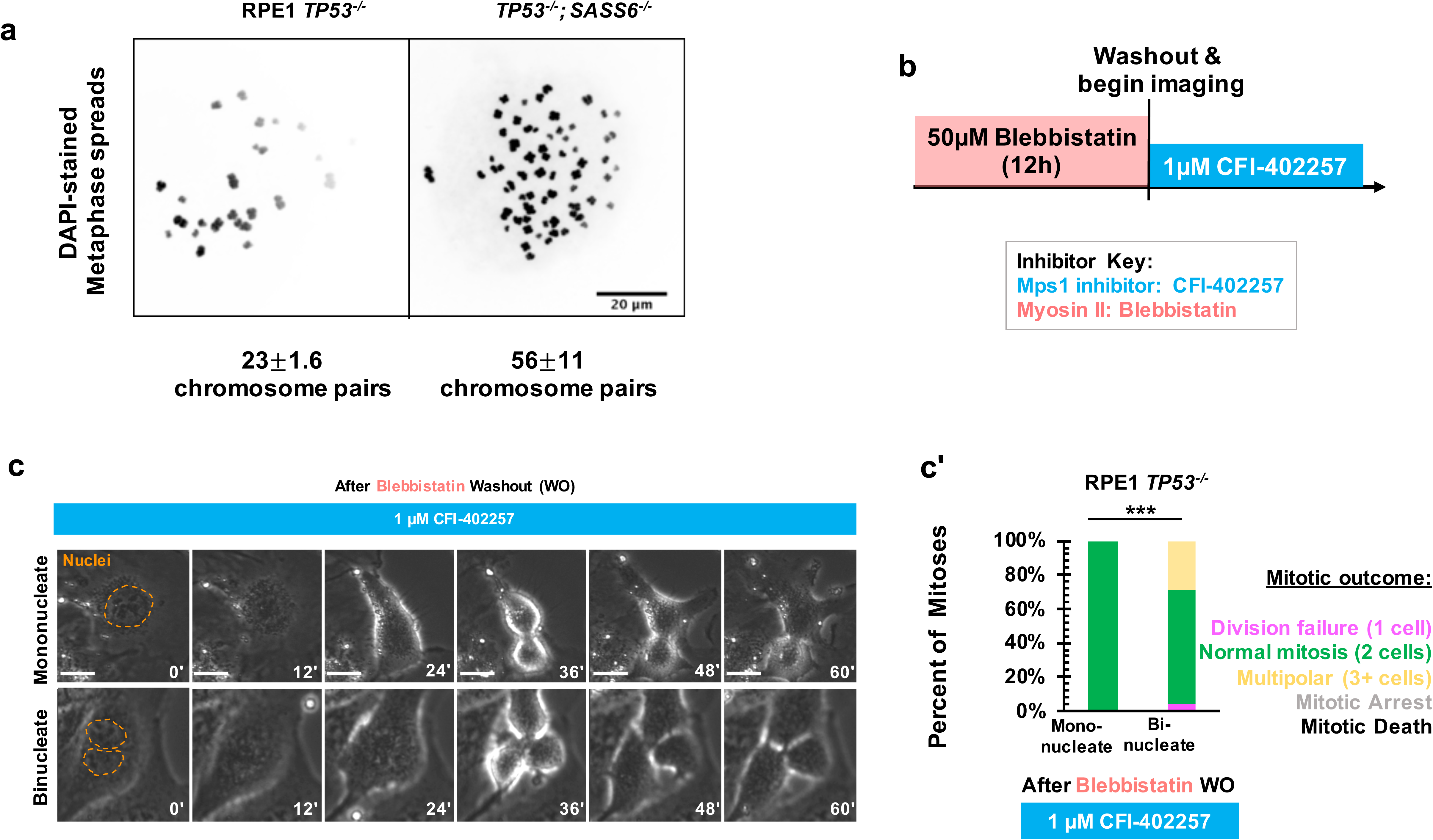
Polyploidy is not sufficient to confer MPS1-inhibition-sensitive division failure. **(a)** Metaphase spreads from RPE1 *TP53^-/-^* or RPE1 *TP53^-/-^; SASS6^-/-^* cells. Indicated beneath example images are the mean and standard errors of chromosome pairs for the two genotypes. Chromosomes were stained with DAPI. **(b)** Schematic of experimental setup. REP1 TP53-/- cells were treated with 50 μM blebbistatin for 12h, washed, and then treated with 1μM CFI-402257 and imaged live via phase microscopy. **(c)** Example mitoses in mononucleate or binucleate cells treated with 1 µM CFI-402257 after washout (WO) of blebbistatin. **(c’)** Quantification of mitotic outcome of RPE1 *TP53^-/-^*or U2OS cells imaged in DMSO, 5 μM STLC, 1 μM CFI-402257, or 5 μM STLC and 1 μM CFI-402257 together. Shown are the percentages for each fate of the total mitotic observations. *n*>48 cells per condition Significance was determined through a Fisher’s exact test. In all cases, not significant (n.s.) denotes *p*>0.05 and *** denotes p<0.001. All scale bars: 10 μm. **Figure3_Source_Data_1.zip: Source data for Figure 3c’**

### Early mitotic MTOC separation is necessary for cell division in the absence of SAC-mediated mitotic delay

Since polyploidy was not sufficient to cause division failure in the absence of the SAC, we next tested whether this phenotype resulted from temporal differences in establishing bipolarity in the absence of centrosomes. This would be consistent with the monopolar spindle seen under MPS1 inhibition in Figure 1e, and with previous work showing delays in bipolar organization of microtubule-minus-end-associated proteins during early mitosis in acentrosomal cells.^24,25,52^ We used acentrosomal and control U2OS *GFP-TUBA1B* cells to directly visualize microtubule organization in early mitosis. While control cells had two tubulin-dense MTOCs visible before NEBD, acentrosomal cells only established two MTOCs well after NEBD and chromosome condensation (Figure 4a-a’).

**Figure 4.**
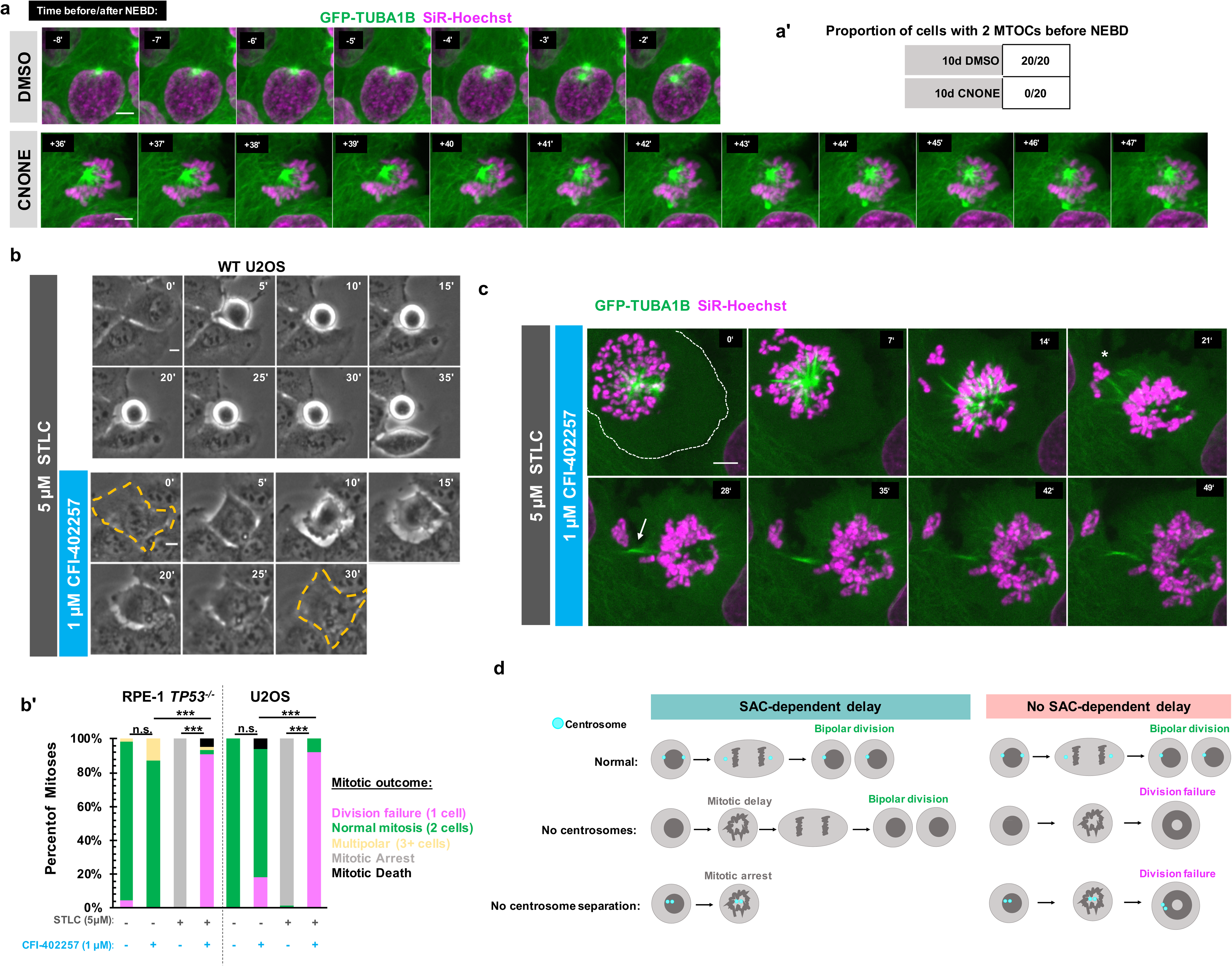
Early mitotic MTOC separation is necessary for cell division in the absence of SAC-mediated mitotic delay. **(a)** Confocal timelapse imaging of U2OS cells pretreated for 10d with either DMSO or centrinone (125 nM). Shown are endogenously tagged α-tubulin (*GFP-TUBA1B*) and DNA (Sir-Hoechst). Time indicates minutes before (-) or after (+) NEBD. **(a’)** Proportion of cells with 2 foci of microtubules before NEBD based on experiments as in (a). **(b)** Live phase imaging of RPE1 *TP53^-/-^* or U2OS cells imaged in 1μM CFI-402257 or 5 μM STLC and 1μM CFI-402257 together. **(b’)** Quantification of mitotic outcome of RPE1 *TP53^-/-^*or U2OS cells imaged in DMSO, 1μM CFI-402257, 5 μM STLC, or 5 μM STLC and 1μM CFI-402257 together. Shown are the percentages for each fate of the total mitotic observations. Significance was determined through a Fisher’s exact test. *n*>50 cells per condition **(c)** Confocal timelapse imaging of U2OS cells imaged in 5μM STLC with 1μM CFI-402257. Shown are endogenously tagged α-tubulin (*GFP-TUBA1B*) and DNA (Sir-Hoechst). Asterisk indicated extruded chromosomes. Arrow indicated midbody-like α-tubulin structure. **(d)** Graphical summary of results. In all cases, not significant (n.s.) denotes *p*>0.05 and *** denotes p<0.001. All scale bars: 10 μm. **Figure4_Source_Data_1.zip: Source data for Figure 4b’**

Given this result, one possible explanation for the robustness of cell division in cells with centrosomes is that having two separate and functional MTOCs at the beginning of mitosis facilitates division in the absence of a SAC-mediated delay. We therefore tested whether a monopolar spindle, such as in acentrosomal cells early in mitosis, is sufficient to cause division failure upon MPS1 inhibition, even in the presence of centrosomes. We treated RPE1 or U2OS cells containing centrosomes with (+)-S-Trityl-L-cysteine (STLC), an inhibitor of the bipolar kinesin Eg5 (also known as Kif11 or kinesin-5), resulting in monopolar mitotic spindles.^53,54^ As expected, treatment with STLC alone caused mitotic arrest (Figure 4b-b’); however, inhibiting MPS1 in addition to Eg5 resulted in exit from mitotic arrest and division failure (Figure 4b-b’) similar to the phenotype observed in acentrosomal cells.

To better understand how centrosome separation in early mitosis allows normal mitosis to occur without MPS1-dependent delay, we imaged Eg5-inhibited cells, with or without MPS1 inhibition, via confocal microscopy. Cells treated with both STLC and CFI-402257 formed monopolar spindles but then exited from mitosis (Figure 4c). Only one cell resulted from this failed division. However, frequently some chromosomes became extruded (Figure 4c, asterisk) during what appeared to be a highly asymmetric division, with both centrosomes remaining unseparated in the primary cell. In addition, a midbody-like structure (Figure 4c, arrow) formed between the main resultant cell and the extruded portion. Taken together, these results show that the early separation of centrosomes prevents monopolar spindles from persisting, allowing a timely completion of mitosis even without the SAC.

Here, we show that acentrosomal cells rely on SAC-mediated delay, not simply to correct chromosome attachment errors, but even to build a bipolar spindle and divide into two daughters (Figure 4d). Because centrosomes form two MTOCs very early in mitosis and can be efficiently separated into two spindle poles by Eg5 activity, they provide a “timely two-ness” to mitotic organization. This rapid establishment of bipolarity normally allows cells to successfully divide in the absence of a SAC-induced delay. The finding of such ‘cross-talk’ between centrosomes and the SAC leads to exciting questions about spindle organization and function. For instance, it remains unknown why acentrosomal cells, whether created in a dish or during female meiosis in many metazoans, are prone to chromosome missegregation despite the ability to assemble bipolar spindles.^31,47,48,55,56^ It is known that SAC activity weakens over time during prolonged arrest, characterized by decreasing levels of cyclin B globally and MAD2 at kinetochores, thus enabling mitotic slippage.^57–59^ Therefore, a fertile path of investigation will be to determine whether the efficiency that centrosomes provide to spindle bipolarity enables important temporal coordination between SAC-mediated delay and correction of spindle defects by ensuring that these corrections occur during a time window when global cyclin B and kinetochore MAD2 levels are still elevated. On a more translational note, existing MPS1 inhibitors are under investigation as chemotherapeutic agents,^60–63^ and the presence of acentrosomal cells in cancers is now being appreciated.^47,64^ Given our results, centrosome presence or loss may be a relevant clinical marker to consider when predicting effects of MPS1 inhibition on division outcome.

## Acknowledgements

Centrinone was a gift of Karen Oegema (University of California, San Diego). hTERT-RPE1 *TP53^-/-^* and *TP53^-/-^;SASS6^-/-^* cells were a gift from Meng-Fu Bryan Tsou (Sloan Kettering Institute). The plasmid AICSDP-4:TUBA1B-mEGFP was a gift from Allen Institute for Cell Science (Addgene plasmid # 87421; RRID:Addgene_87421). The plasmid plentiCRISPRv2 was a gift from Feng Zhang (Addgene plasmid # 52961 ; RRID:Addgene_52961). We thank Alexandra F. Long (Stanford University), Ljiljana Milenkovic (Stanford University), and Sabrina Ergun (Princeton University) for critical feedback on the manuscript. K.F. was supported by the National Institutes of Health (NIH) under award number T32GM007276. J.T.W. was supported by the National Institute of General Medical Sciences of the NIH under award number K99GM131024. This work was supported by the NIH under grant R35GM130286 to T.S. The content is solely the responsibility of the authors and does not necessarily represent the official views of the NIH.

## Materials and Methods

### Cell Lines and Culture

hTERT RPE-1; *TP53^-/-^* and hTERT RPE-1; *TP53^-/-^; SASS6^-/-^* were a gift from Bryan Tsou and described previously.^32,65^ U2OS cells were acquired from the ATCC. hTERT RPE-1 cells of all genotypes were cultured in DMEM/F-12 (with L-glutamine and 15 mM HEPES, Corning) supplemented with 10% fetal bovine serum (FBS; Gemini Bio-Products), and 1% Pen-Strep (final concentrations: 100 U/mL penicillin G and 0.1 mg/mL streptomycin, Gemini Bio-Products). U2OS cells were cultured in DMEM (with L-glutamine, 4.5g/L glucose and sodium pyruvate, Corning) supplemented with 10% CCS and 1% Pen-Strep. All cells were cultured at 37 °C with 5% CO2. Cells were routinely tested for mycoplasma with MycoAlert Mycoplasma Detection Kit (Lonza) as per manufacturer instructions.

### Drugs and Live Stains

Drugs were used the indicated concentrations in figures and figure legends and came from the following sources: CFI-402257 (Caymen Chemical Company, stock concentration: 4mM in DMSO), proTAME (R&D Systems, stock concentration: 20mM in DMSO), S-Trityl-L-cysteine (STLC) (Sigma-Aldrich, stock concentration: 10mM in DMSO); MPS1 IN-1 (Caymen Chemical Company, stock concentration: 10 mM in DMSO); AZD1152-HQPA (also called Barasertib-HQPA, Caymen Chemical Company, stock concentration: 50mM in DMSO); (S)-(-)-Blebbistatin (Toronto Research Chemicals, stock concentration 100mM in DMSO). Centrinone, a generous gift from Karen Oegema, was always used at 125nM (stock concentration: 125μM in DMSO). For live imaging, SiR-DNA/SiR-Hoechst (Spirochrome) was used at 50 nM (stock concentration: 5mM in DMSO).

### Live phase imaging

Cells were plated 5x10^4^ cells/well in glass bottom 12-well plates (No. 1.5 Coverslip, 14 mm Glass Diameter, Uncoated 12 well plate, Mattek) or 24-well plated (No. 1.5 Coverslip, 14 mm Glass Diameter, Uncoated 24 well plate, Mattek) to facilitate simultaneous imaging of various drug concentrations. Phase images were acquired with a Keyence microscope with either a Nikon 20x or a Nikon 10x objective. Temperature was maintained at 37°C in a humidified chamber with 5% CO_2_ flowed in. Images were acquired every 4 or 5 min depending on number of positions acquired.

### Confocal Microscopy

For both fixed and live cells, samples were imaged with an Zeiss Axio Observer microscope (Carl Zeiss) with a confocal spinning-disk head (Yokogawa Electric Corporation, Tokyo, Japan), PlanApoChromat 63×/1.4 NA objective, and a Cascade II:512 electron-multiplying (EM) CCD camera (Photometrics, Tucson, AZ) or PRIME: BSI backside illuminated CMOS camera run with SlideBook 6 software (3i, Denver, CO). Excitation lasers were 405nm, 488nm, 661nm, and 640nm.

### CRISPR endogenous tagging of *TUBA1B* with mEGFP in U2OS cells

To create U2OS cells expressing mEGFP-TUBA1B, we followed the same CRISPR/Cas9 gene editing design as in Roberts et al.^41^ Briefly, U2OS cells were co-transfected with pLentiCRISPRv2^66^ (Addgene plasmid # 52961 ; RRID:Addgene_52961) expressing a gRNA targeted against *hTUBA1B* (gGATGCACTCACGCTGCGGGA) and an mEGFP plasmid with 1 kb homology arms (AICSDP-4:TUBA1B-mEGFP, Allen Institute, Addgene plasmid # 87421; RRID:Addgene_87421)^41^ mutated for the gRNA binding site. Two days after transfection, cells were split into a fresh plate with 1 μg/mL puromycin. After two days of selection, cells were placed into fresh media. Single cell clones were obtained by serial dilution, and clones were screened by PCR with the following primers: forward primer 5’-GGGGTGCTGGGTAAATGGAG reverse primer 5’-CGGTTTAGGATGGGAAGGTAACATT.

Clones were then assessed via live imaging for GFP fluorescence, Western blot for GFP tagging of α-tubulin, the ability of this tagged α-tubulin to be acetylated, and the absence of Cas9 protein in the established clone, and immunofluorescence for localization of GFP-TUBA1B to spindle microtubules. The chosen clone was sequenced and found to have one allele edited with the expected insertion and one allele unedited at the *TUBA1B* locus.

### Live fluorescent imaging of microtubules and DNA

U2OS *GFP-TUBA1B* cells were plated into 35 mm glass bottom dishes (Fluorodish) in normal grown medium. 30 min before onset of imaging, medium was changed to phenol free DMEM (with L-glutamine, 4.5g/L glucose and sodium pyruvate, Corning) containing 10% CCS (Gemini bio-Products), 50nM SiR-Hoechst (Spirochrome), and any drug treatments (STLC or CFI-402257). Temperature was maintained at 37 °C, and the imaging dish was kept in a chamber into which humidified 5% CO2 was flowed in. Images were acquired every minute via confocal microscopy (488nm and 640nm laser lines).

### Antibodies and Fixed Stains

Primary antibodies used for immunofluorescence (IF) and Western blot (WB): mouse IgG1 anti-a-tubulin, clone DM1A (1:1,000IF, 1:5,000WB, Sigma-Aldrich T6199; RRID:AB_477583); mouse IgG2_b_ anti-centrin3, clone 3e6 (1:1,000 IF, Novus Biological H00001070-M01, RRID:AB_537701); mouse IgG1anti-y-tubulin, clone GTU-88 (1:1000 IF, Sigma-Aldrich T5326; RRID:AB_477584); rabbit anti-CP110 (1:200 IF, Proteintech 12780-1-AP; RRID:AB_10638480); mouse IgG2_b_ anti-Lys-40-acetylated a-tubulin, clone 6-11B-1 (1:1,000 IF, 1:10,000 WB, Sigma-Aldrich T6793; RRID:AB_477585); mouse IgG2_a_ anti-GFP, clone 3e6 (1:1,000 IF, 1:5000 WB, Thermo Fischer A-11120; RRID:AB_221568); mouse IgG1 anti-Cas9, clone 7A9 (1:1000 WB, BioLegend 698301, RRID: AB_2715788). Secondary antibodies conjugated to AlexaFluor (Thermo Fisher/Invitrogen) were used for IF. Secondary antibodies conjugated to IR dyes (BioRad) were used for Western blots. REVERT total protein stain (BioRad) was used as indicated by manufacturer.

### Immunofluorescence

#1.5 coverslips were washed with 70% Ethanol and dried under UV light before use. Cells were fixed for 7 min in 100% -20°C methanol. Cells were washed 3x at room temperature (RT) with PBS and blocked for 10 min with PBS-BT (PBS + 3% BSA + 0.1% Triton-X). Primary antibodies diluted in blocking solution were incubated for 1h at RT. Cells were then washed 3x 10 min in blocking solution and then incubated with secondary antibodies, also in blocking solution, for 45 min at RT. Cells were washed 2x 10 min with blocking solution, and 1x 10 min with PBS (containing 5 μg/ml DAPI (4′,6-diamidino-2-phenylindole), if used). Samples were then mounted in Mowiol mounting medium (Polysciences) in glycerol containing 2.5% 1,4-diazabicyclo-(2,2,2)-octane (DABCO, Sigma-Aldrich) antifade.

### Metapahse spreads

Cells were incubated for 3h in 10 ng/μL nocodazole in normal growth medium. Cells were then trypsanized and pelleted and resuspended in ice cold 0.56% KCl solution, incubated at RT for 7 min for hypotonic lysis to occur, then pelleted. Pellets were then disrupted and cells were resuspended and fixed in fixative (75% v/v methanol; 25% v/v glacial acetic acid) added dropwise. Cells were pelleted and resuspended in fixative. 20μL drops were placed onto ethanol-cleaned slides, which were left to dry. Spreads were then overlayed by Mowiol mounting medium (Polysciences) in glycerol containing 2.5% 1,4-diazabicyclo-(2,2,2)-octane (DABCO, Sigma-Aldrich) antifade containing 5 μg/ml DAPI (4′,6-diamidino-2-phenylindole). #1.5 coverslips were placed on top, and samples were imaged via confocal microscopy.

### SDS-PAGE and Western blot

Samples were separated by 12% SDS-PAGE reducing gels (5% stacking gels) and transferred to nitrocellulose membrane at 4°C. Blots were blocked in 5% (w/v) non-fat dry milk in TBST (TBS + 1% Tween-20). Primary and secondary antibody staining were performed in blocking solution. When used, total protein on nitrocellulose membranes was stained via REVERT Total Protein (LiCOR) as per manufacturer instructions and imaged before immunodetection. REVERT-stained blots and immunoblots were imaged using a LiCor Odyssey.

### Statistical analysis

For categorical data, significance was determined a Fisher’s exact test. For quantitative data, significance was determined via Welch’s *t*-test. The test performed is also specified in the figure legend. In all cases, not significant (n.s.) denotes *p*>0.05; * denotes p<0.05; ** denotes p<0.01; *** denotes p<0.001. Statistical tests were performed in R Studio. SEM was calculated in Excel or R-Studio as one standard deviation divided by the square-root of the *n*. The given *n* in figure legends refers to biological replicates from two or more technical replicates. Sample size was determined using the maximum number of cells practically observable given constraints of the experiments. For exact *n*, please consult the source data spreadsheet.

### Novel Materials Availability Statement

The newly created U2OS *mEGFP-TUBA1B* cell line generated in this work is available through contacting Tim Stearns (Rockefeller University) and/or Jennifer T. Wang (Department of Biology, Washington University in St. Louis).

---

**Figure 1-supplement 1. Treatment with alternate MPS1 inhibitor MPS1 IN-1 phenocopies treatment with CFI-402257. (a)** Quantification of daughter number of cells of the given pre-treatment imaged in DMSO or 10 μM MPS1 IN-1. Shown are the percentages for each fate of the total mitotic observations. Significance was determined through a Fisher’s exact test. *n*≥41 cells per condition. **(b)** Schematic of inhibitors and targets. In all cases, not significant (n.s.) denotes *p*>0.05 and *** denotes p<0.001. All scale bars: 10 μm.

**Figure1_Supp1a_Source_Data_1.zip: Source data for Figure 1-supplement 1a**

**Figure 1-supplement 2. MPS1 inhibition results in abnormal nuclear morphology, but not binuclearity, in acentrosomal cells. (a)** DAPI staining of cells after 72 h treatment with DMSO or 1 μM CFI-403357. (**a’**) Shown are the percentages of cells with each indicated nuclear morphology for interphase nuclei. *n*≥100 cells per condition. In all cases, not significant (n.s.) denotes *p*>0.05, * denotes p<0.05, **denotes p<0.01, and *** denotes p<0.001.

**Figure1_Supp2a_Source_Data_1.zip: Source data for Figure 1-supplement 2a**

**Figure 1-supplement 3. Creation and validation of *GFP-TUBA1B* U2OS cells. (a)** Agarose DNA gel of PCR genotyping of unedited cells, and edited pool of cells, and the edited clone of cells (clone 1-F3) used in future experiments. PCR primers in material and methods. **(b)** Western blot of α-tubulin and GFP levels in unedited cells (transfected with Cas9 plasmid but not gRNAs), unedited cells, and edited cells. **(c)** Western blot of Cas9 levels in cells freshly transfected with Cas9, unedited cells, and edited cells. This confirms that the Cas9 was transient and is no longer present in the edited cells and had been expressed only transiently. **(d)** Western blot of α-tubulin and acetylated-α-tubulin levels in unedited and edited cells, showing that the GFP-tagged α-tubulin can be acetylated (arrow). **(e)** Immunofluorescence of GFP (green), all α-tubulin (magenta) in U2OS *GFP-TUBA1B* cells in prometaphase (top) or metaphase (bottom). Scale bars: 10 μm.

**Figure1_Supp3_Source_Data_1.zip: Source data for Figure 1-supplement 3b, c, d**

**Figure 1-supplement 4. Division failure in acentrosomal cells due to Mps1 inhibition is reversible after centrosome return. (a)** Immunofluorescence staining of RPE-1 *TP53^-/-^* or U2OS cells treated with DMSO or 125 nM centrinone (CNONE) for 10 days and then washed out for 10 days. DAPI is shown in blue, acetylated-α-tubulin in green, and CP110 in magenta. **(a’)** Quantification of a. Graphed are means and S.E.M. Significance was determined through a Fisher’s exact test. *n*≥ 100 cells per condition. **(b)** Quantification of mitotic duration in cells of indicated genotype after 10 d DMSO followed by 10 d washout (grey) or 10d centrinone (125 nM) followed by 10 d washout (blue). Points represent individual cells; boxplots represent mean and interquartile range. Significance was determined through Welch’s *t*-test. *n*≥ 25 cells per condition. **(c)** Quantification of daughter number of cells of the given pre-treatment imaged in DMSO or CFI-402257 (1 μM). Shown are the percentages for each fate of the total mitotic observations. Significance was determined through a Fisher’s exact test. *n*≥40 cells per condition. In all cases, not significant (n.s.) denotes *p*>0.05 and *** denotes p<0.001. All scale bars: 10 μm.

**Figure1_Supp4_Source_Data_1.zip: Source data for Figure 1-supplement 4a’, b, c**

