## supplemental figures for "Spindle assembly checkpoint-dependent mitotic delay is required for cell division in absence of centrosomes"

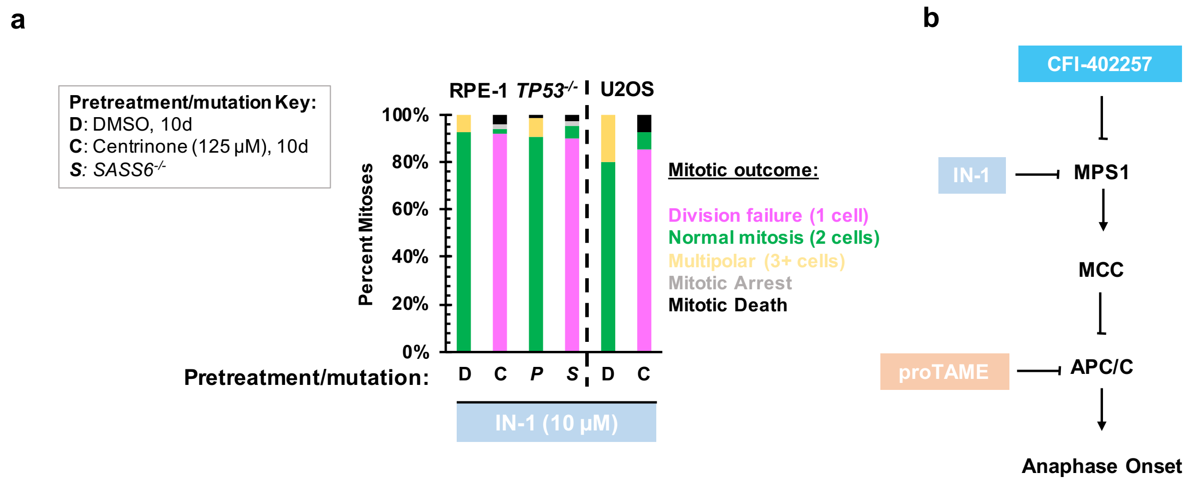


**Figure S1. Treatment with alternate MPS1 inhibitor MPS1 IN-1 phenocopies treatment with CFI-402257. (a)** Quantification of daughter number of cells of the given pre-treatment imaged in DMSO or 10 µM MPS1 IN-1. Shown are the percentages for each fate of the total mitotic observations. Significance was determined through a Fisher's exact test. $n$=50 cells per condition. **(b)** Schematic of inhibitors and targets. In all cases, not significant (n.s.) denotes $p$>0.05 and *** denotes p<0.001. All scale bars: 10 µm.

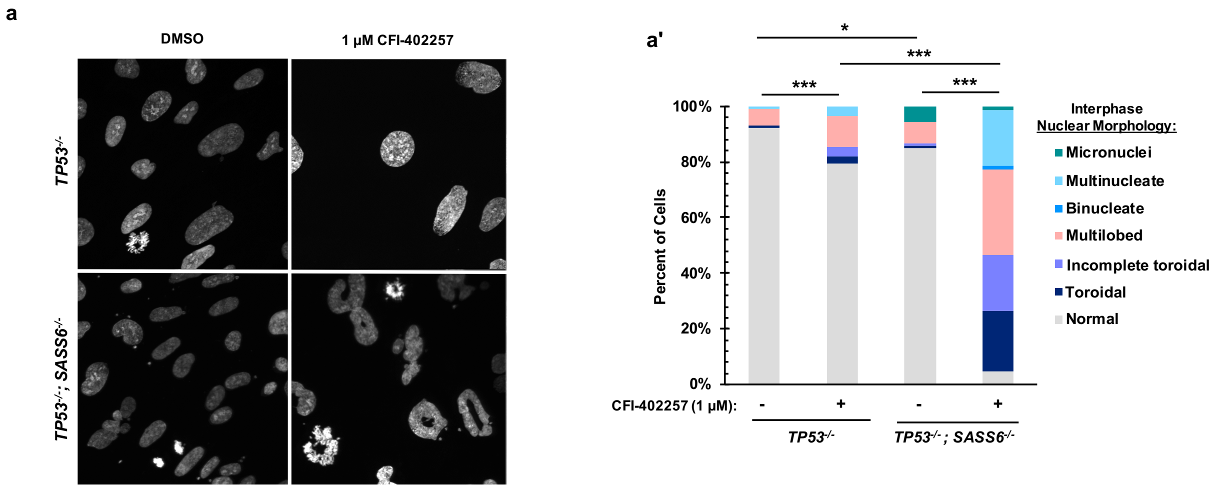


**Figure S2. MPS1 treatment results in abnormal nuclear morphology, but not binuclearity, in acentrosomal cells. (a)** DAPI staining of cells after 72 h treatment with DMSO or 1 µM CFI-403357. (**a'**) Shown are the percentages of cells with each indicated nuclear morphology for interphase nuclei. *n*=100 cells per condition. In all cases, not significant (n.s.) denotes *p*>0.05, * denotes p<0.05, **denotes p<0.01, and *** denotes p<0.001.

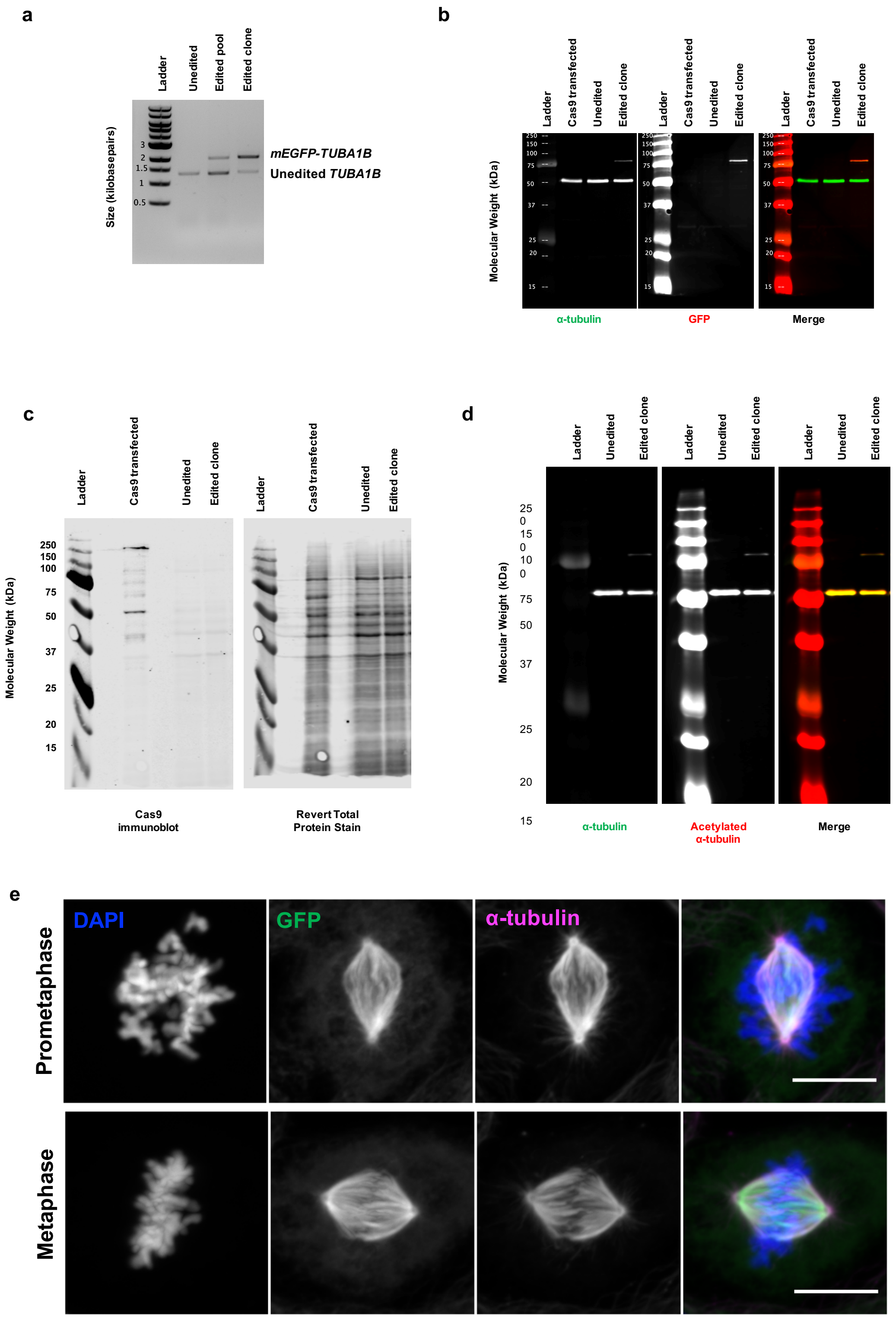


**Figure S3. Creation and validation of *GFP-TUBA1B* U2OS cells. (a)** Agarose DNA gel of PCR genotyping of unedited cells, and edited pool of cells, and the edited clone of cells (clone 1-F3) used in future experiments. PCR primers in material and methods. **(b)** Western blot of α-tubulin and GFP levels in unedited cells (transfected with Cas9 plasmid but not gRNAs), unedited cells, and edited cells. **(c)** Western blot of Cas9 levels in cells freshly transfected with Cas9, unedited cells, and edited cells. This confirms that the Cas9 was transient and is no longer present in the edited cells and had been expressed only transiently. **(d)** Western blot of α-tubulin and acetylated-α-tubulin levels in unedited and edited cells, showing that the GFP-tagged α-tubulin can be acetylated (arrow). **(e)** Immunofluorescence of GFP (green), all α-tubulin (magenta) in U2OS *GFP-TUBA1B* cells in prometaphase (top) or metaphase (bottom). Scale bars: 10 μm.

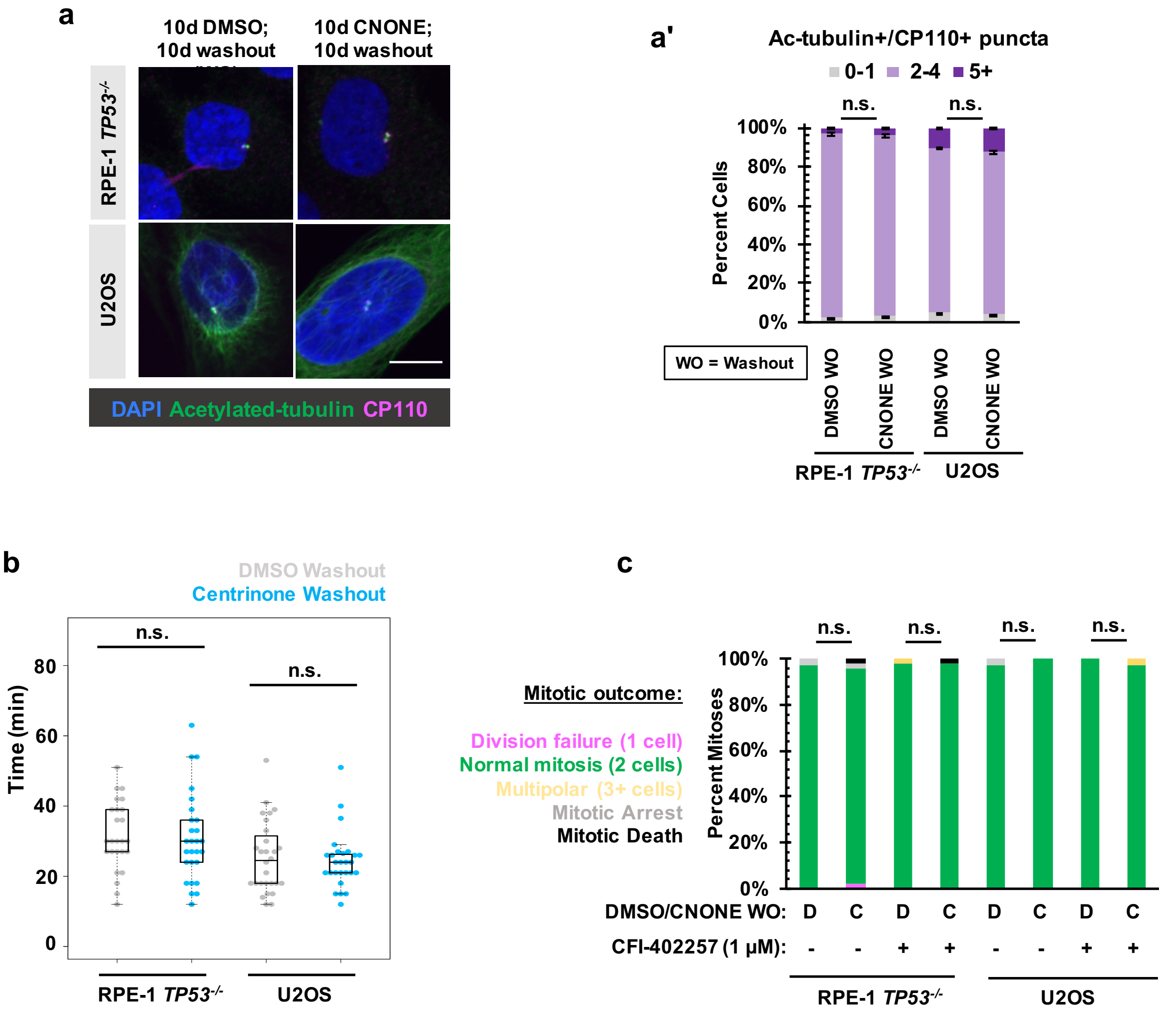


**Figure S4. Division failure in acentrosomal cells due to Mps1 inhibition is reversible after centrosome return.** **(a)** Immunofluorescence staining of RPE-1 *TP53*$^{-/-}$ or U2OS cells treated with DMSO or 125 nM centrinone (CNONE) for 10 days and then washed out for 10 days. DAPI is shown in blue, acetylated-α-tubulin in green, and CP110 in magenta. **(a')** Quantification of c. Graphed are means and S.E.M. Significance was determined through a Fisher's exact test. **(b)** Quantification of mitotic duration in cells of indicated genotype after 10 d DMSO followed by 10 d washout (grey) or 10d centrinone (125 nM) followed by 10 d washout (blue). Points represent individual cells; boxplots represent mean and interquartile range. Significance was determined through Welch's *t*-test. **(c)** Quantification of daughter number of cells of the given pre-treatment imaged in DMSO or CFI-402257 (1 µM). Shown are the percentages for each fate of the total mitotic observations. Significance was determined through a Fisher's exact test. In all cases, not significant (n.s.) denotes $p>0.05$ and *** denotes p<0.001. All scale bars: 10 µm.
